# The SHAPE of logistic growth shows that serial passaging biases fixation probability

**DOI:** 10.1101/392944

**Authors:** Jonathan Dench

**Affiliations:** Department of Biology, University of Ottawa, Ottawa, ON, Canada

## Abstract

The forward time simulation tool rSHAPE (R-package for Simulated Haploid Asexual Population Evolution) was designed to complement the theoretical and empirical study of evolution. Included with rSHAPE are functions to programmatically build, run, and initially process results of an evolutionary experiment defined by the range of experimental conditions. As experimental evolution often studies both the emergence and fate of *de novo* mutants, I validated rSHAPE by confirming its ability to replicate seminal theoretical expectations concerning changes in fitness through time and the fixation probability of mutants. As an example of how rSHAPE can support both theoretical and empirical research, I applied rSHAPE to study how the laboratory protocol of serial passaging, common in microbial experimental evolution, affects the fixation probability of *de novo* mutants. Unlike related theoretical work which modelled growth as effectively exponential (Wahl *et al.*, 2002), this study considered populations experiencing logistic growth which is common to microbes undergoing serial passaging. In contrast to exponential growth, when a population undergoes logistic growth the probability of a mutant arising and eventually fixing depends upon when a mutant arises during a growth phase. Users can download software and documentation for rSHAPE through CRAN at https://cran.r-project.org/web/packages/rSHAPE/index.html, or via GitHub at https://www.github.com/Jdench/SHAPE_library/.

## Introduction

While certain independent evolutionary drivers have been elucidated, such as selective context or large effect beneficial mutations, researchers are increasingly interested in studying the fate of populations, and the mutations they carry, in a more wholistic context (Losos *et al.*, 2013). Though experimental evolution can be paired with next generation sequencing technologies in order to track the emergence and fate of mutations (Goodwin *et al.*, 2016), these *in vivo/in vitro* approaches are limited by availability of physical or financial resources. Even when experimental design is optimised, it is still not commonly feasible for researchers to fully sequence whole genomes, of whole populations, at multiple time intervals, for each replicate of an experiment. Such an approach is not fanciful idealism as the interaction among mutations can affect evolutionary outcome (Weinreich *et al.*, 2006; Phillips, 2008; Domingo *et al.*, 2019). Furthermore, at least one well funded research group has sequenced whole replicate populations, at multiple time points, shedding new light on evolutionary dynamics (Good *et al.*, 2017). While development of evolutionary theory with, and analytical analysis of, existing data are much less costly approaches, we need to combine both evolutionary theory and empirical study to improve our understanding of evolution in increasingly complex systems (de Visser and Krug, 2014). Simulations, or *in silico* evolution, can leverage theoretical models based on empirical evidence to support both theoretical and empirical study of evolutionary biology, while being limited only by the available software and computational resources.

Simulation software complement theoretical and empirical study by allowing the comparison of empirical observation to the dynamics expected from various theoretical models and combinations of evolutionary parameters. There exist simulation tools to track changes in population demographics (Guillaume and Rougemont, 2006) or specific mutations (Lambert *et al.*, 2008; Dalquen *et al.*, 2012; Zanini and Neher, 2012; Arnold *et al.*, 2018), though most of these implement a single theoretical model and provide only limited output. The AVIDA(Beckmann *et al.*, 2010) and AEVOL(Knibbe *et al.*, 2007) tools are agent-based models for studying genetic changes but their evolutionary systems are genomic abstractions that are practical for their implementation but make it unclear to what extent their results apply to living systems. A powerful tool for simulating microbial experimental evolution is the Haploid Evolutionary Constructor (HEC) (Lashin *et al.*, 2014), though it simplifies evolution of the genome by mapping each locus to a single quantitative trait value and assumes mutations do not interact. Knowing no single framework to simulate experimental evolution while tracking detailed changes in both population demographics and all mutations in large genomes (*i.e.* millions of mutational sites), I found the existing software impractical, or not applicable, for comparing evolution across different models of growth and fitness landscape scenarios. To address this, I developed the R-package for SHAPE (Simulated Haploid Asexual Population Evolution - rSHAPE).

The simulation tool rSHAPE was designed to simulate microbial experimental evolution in a framework that can readily be expanded to different models, and evolutionary scenarios, thanks to a modular programming design. I chose to simulate microbial evolution experiments as they are commonly used for the manipulative study of evolution. These experiments often study how the selective context influences the emergence and persistence of *de novo* mutants arising from an initially isogenic population, but are also used to estimate underlying evolutionary parameters such as the fitness landscape of selective contexts. The design of many microbial evolution experiments requires the practice of serial passaging, a protocol in which a small fraction of stationary phase communities is transferred to fresh growth media to extend the evolutionary time of experiments. When communities are well mixed prior to serial passaging, loss of mutants is proportional to their representation in the community. While serial passaging is expected to result in the loss of rare mutants, theory suggests that the probability of a mutant arising during exponential growth, and subsequently surviving many rounds of serial passaging, is roughly uniform (Wahl *et al.*, 2002). For experimental microbial populations, continuous cell division (similar to exponential growth conditions) will only generally be achieved in chemostats. Many microbial experiments avoid chemostats due to their expense and the difficulty with both setup and maintenance. Thus, most experimental microbial communities evolve under logistic growth conditions punctuated by population bottlenecks (*i.e.* serial passaging). It remains unclear how serial passaging may affect the joint probability of mutants arising and subsequently fixing in evolving populations.

In this paper I begin by explaining the framework of rSHAPE. I then report on rSHAPE’s ability to reproduce expected trends in fitness through time as well as the probability of a *de novo* mutant fixing under the conditions of several theoretical models. I then provide one example of how rSHAPE can be used to study evolutionary scenarios outside the scope of existing analytical models. I used rSHAPE to test if serial passaging influences the fixation probability of a *de novo* mutant arising in a population that undergoes logistic growth.

## Design and implementation

### Simulating evolution

The software rSHAPE is a forward time, discrete step, *in silico* experimental evolution system that tracks changes in population demographics. Populations are composed of haploid genotypes that are tracked by which of their *L* binary state sites (*i.e.* genome of length *L*) carry mutations and where a value of 0 represents the unmutated state and 1 depicts mutation. Note that while it may be practical to consider each site as a nucleotide base, each site could equally be considered as any mutable element which influences genotype fitness (*e.g.* a codon or gene). To define the experimental conditions, rSHAPE includes the function *defineSHAPE* which allows users to directly set 30 experimental parameters (Appendix **??**, Table **??**). Once the parameters are defined, the experiment is started by calling the function *runSHAPE* which will sequentially simulate the evolution of *n* identical but independent starting populations. In each time step, the population may experience stochastic loss events (*e.g.* a population bottleneck) and each individual within the population may die, reproduce, and/or mutate. Though it may be practical to think of a time step as a biological generation in which each individual of average fitness will produce *r* offspring (where *r* is the growth rate), this need not be the case (*e.g.* users can set the average probability of birth such that *P*_*birth*_ ∈ [0, 1]). Regardless, after each time step rSHAPE records the evolving populations’ demographics as the current number of individuals for each genotype and their numbers of deaths, births, as well as mutants generated including the “parent” genotype from which they arose. By tracking a mutant’s “parent” genotype, the entire evolutionary history can be reconstructed permitting detailed evolutionary analyses. To further support analyses, rSHAPE tracks the full fitness landscape of all explored mutational space. When a “parent” individual experiences a mutation, the explored mutational space represents all possible mutant offspring from which the offspring is eventually selected. At the onset of an rSHAPE experiment, the evolving population will consist entirely of wild-type (0 state for all genomic positions) individuals. In its current form, rSHAPE resembles many microbial evolution experiments in that populations are finite and evolve in a single, well mixed (*i.e.* homogeneous) environment. While these are conditions at the launch of rSHAPE, the starting population, evolutionary environment, as well as other implemented evolutionary models (*e.g.* growth forms, fitness landscapes) could readily be changed due rSHAPE’s modular programming design. This modular design is achieved through rSHAPE being an iterative implementation of functions to calculate the outcome of stochastic events, deaths, births, and mutation (Fig. 1). For a detailed description of the iterative functions called during a run of rSHAPE, please refer to Appendix **??**.

**Figure 1.**
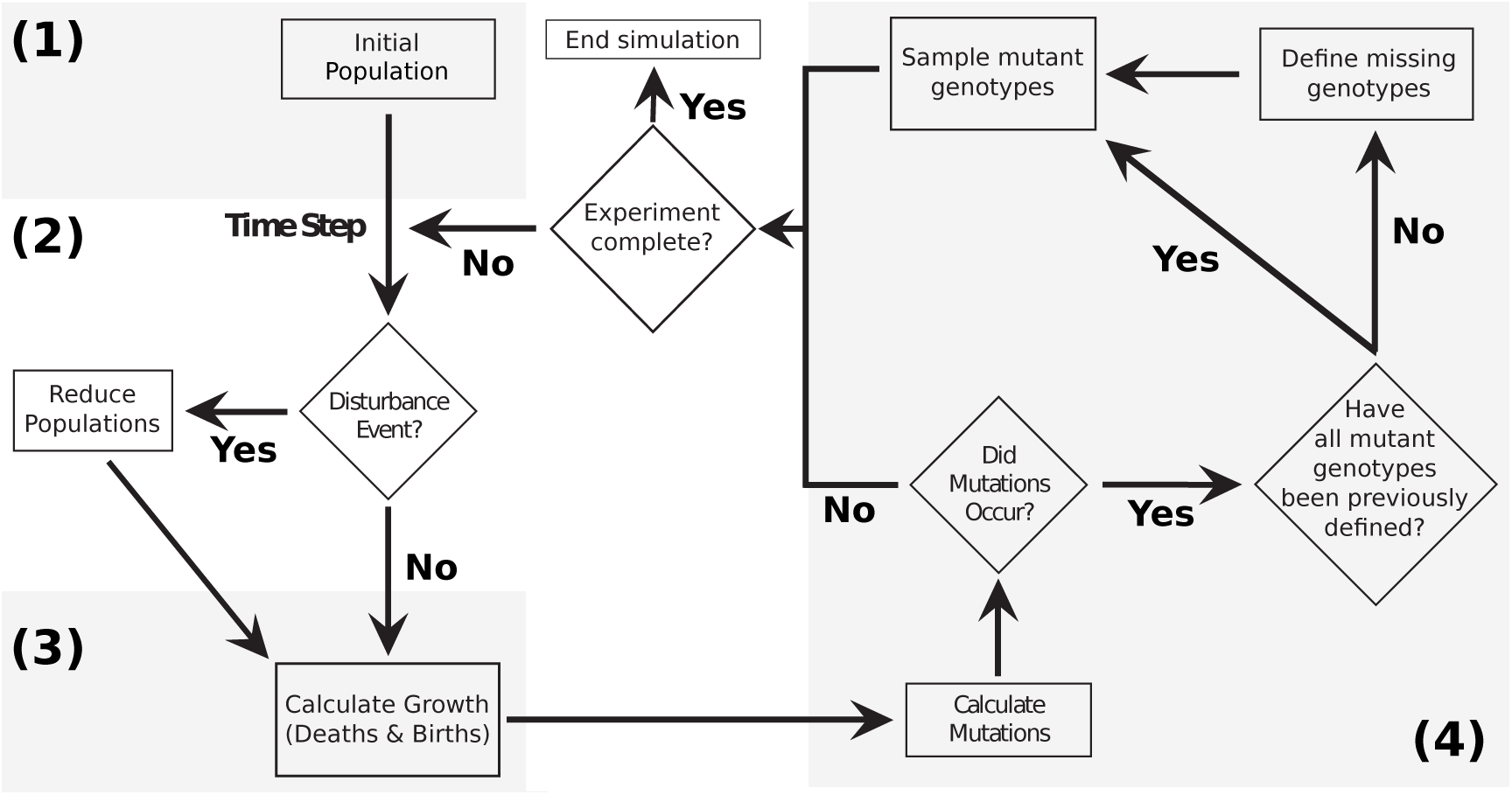
rSHAPE’s flowchart. After an experiment begins, rSHAPE will cycle through discrete time steps which calculate 2) disturbance events, 3) growth (*i.e.* deaths and births), and 4) mutations. Experiments will continue for the pre-determined number of time steps or until there are no individuals living in the population. At the end of the experiment, rSHAPE logs will report the changes occurring in each step and the fitness value of all mutants that could have been produced. The record of mutant fitness values is referred to each time mutation occurs and existing genotype values are retrieved while undefined mutant fitness values are computed and added to the record.

### Running experiments

While populations simulated with rSHAPE are finite, I have set no upper bound on either their size or their genome length. It is possible to define an evolving community of individuals with genomes containing trillions of positions and which grows exponentially without death. It is not practicable - if at all possible - to store in active memory the full demographic changes and genotype fitness values for realistic microbial populations containing billions of individuals with genomes containing millions of sites. To mitigate this challenge, rSHAPE exports the full demographic breakdown, as well as the explored fitness landscape of all genotypes, to SQL databases which act as both the experimental record and a reference when needed during the run. As a result, *in silico* evolution with rSHAPE is less limited by RAM than by disk space and the number of available processors to run independent replicates and parameterisations.

To facilitate rSHAPE’s use for simulating a breadth of experimental conditions, I have included the *shapeExperiment* function that uses template files found here: https://github.com/JDench/SHAPE_library/tree/master/SHAPE_templates. With *shapeExperiment*, rSHAPE builds parameter combinations from the range of values (for each parameter) recorded in the control files. Using these parameter combinations, rSHAPE creates an experimental folder within which all files necessary to run the experiment will be written including automated scripts for running the whole experiment. Before ending each independently replicated simulation, rSHAPE writes a summary log to facilitate analysis. Further, once all simulations in an experiment are complete, users can use the *summariseExperiment* function to combine information from all summary logs and calculate additional summary statistics. Experiments built with *shapeExperiment* will have a script to run *summariseExperiment*. At present, *summarise-Experiment* will create output to facilitate comparison of population dynamics, evolutionary trajectories, and the repeatability of evolution. Future improvements will expand the breadth of summary analytics and allow users to more easily select from among the set of possible summary output.

## Results and discussion

### Validating implementation

To validate the implementation of rSHAPE, I assessed its ability to replicate evolutionary trends expected under various parameterisations of microbial experimental evolution. Since the goal of many microbial evolution experiments is to analyse factors that affect the emergence and maintenance of mutations, I first tested if populations evolving in rSHAPE showed expected fitness trends, and second that the evolutionary dynamics simulated with rSHAPE can replicate the theoretical fixation probability of *de novo* mutants under different growth conditions.

To test rSHAPE’s ability to replicate standard evolutionary trends, I initially tested that the fitness of evolving populations increase through time and that population bottlenecks alter the rate of fitness increase (Wahl *et al.*, 2002) (Fig. 2). For this, I simulated large communities (carrying capacity *K* = 1,000,000) of individuals with 100 genomic positions (*L* = 100), evolving for *T* = 10,000 generations, for each of three different regular bottleneck sizes (*i.e.* disturbance event, dilution factor; *D* ∈ {10, 100, 1000}), where mutant genotype fitness was calculated with either an additive model (a smooth fitness landscape defined by the sum of independent mutational fitness contributions - *i.e.* no epistasis; no interaction between mutations) or a House of Cards (HoC) fitness landscape (Kingman, 1978; Kauffman and Levin, 1987) (the fitness of every genotype is a random value independently drawn from the same distribution - *i.e.* a maximally rugged landscape). The independent effect of all mutations was assumed to be beneficial and so based on extreme value theory (Gillespie, 1984), the random component of all genotype fitness calculations was drawn from an exponential distribution (with rate parameter *λ* = 100). For statistical comparison, I replicated each parameter combination 500 times by initialising five independent fitness landscapes within which evolution was replicated 100 times. For the additive fitness landscape models, I expected fitness to increase until all mutations had fixed since all mutational effects were beneficial. However, each genotype simulated with a HoC fitness landscape is equally likely to be the most fit genotype (*i.e.* global optimum) and many genotypes will be the highest among their nearest mutant “neighbours” (*i.e.* local optimum) (Kauffman and Levin, 1987), so that fitness increase should generally stop long before all mutations fix. From analysis of the evolutionary trends in these first simulations, I found for the additive fitness landscape models that both fitness and the number of mutations increased throughout because not all *L* (100) mutations had yet fixed, whereas in the HoC models evolution stopped after only a few mutations had fixed and a local optima had been found (Fig. 3).

**Figure 2.**
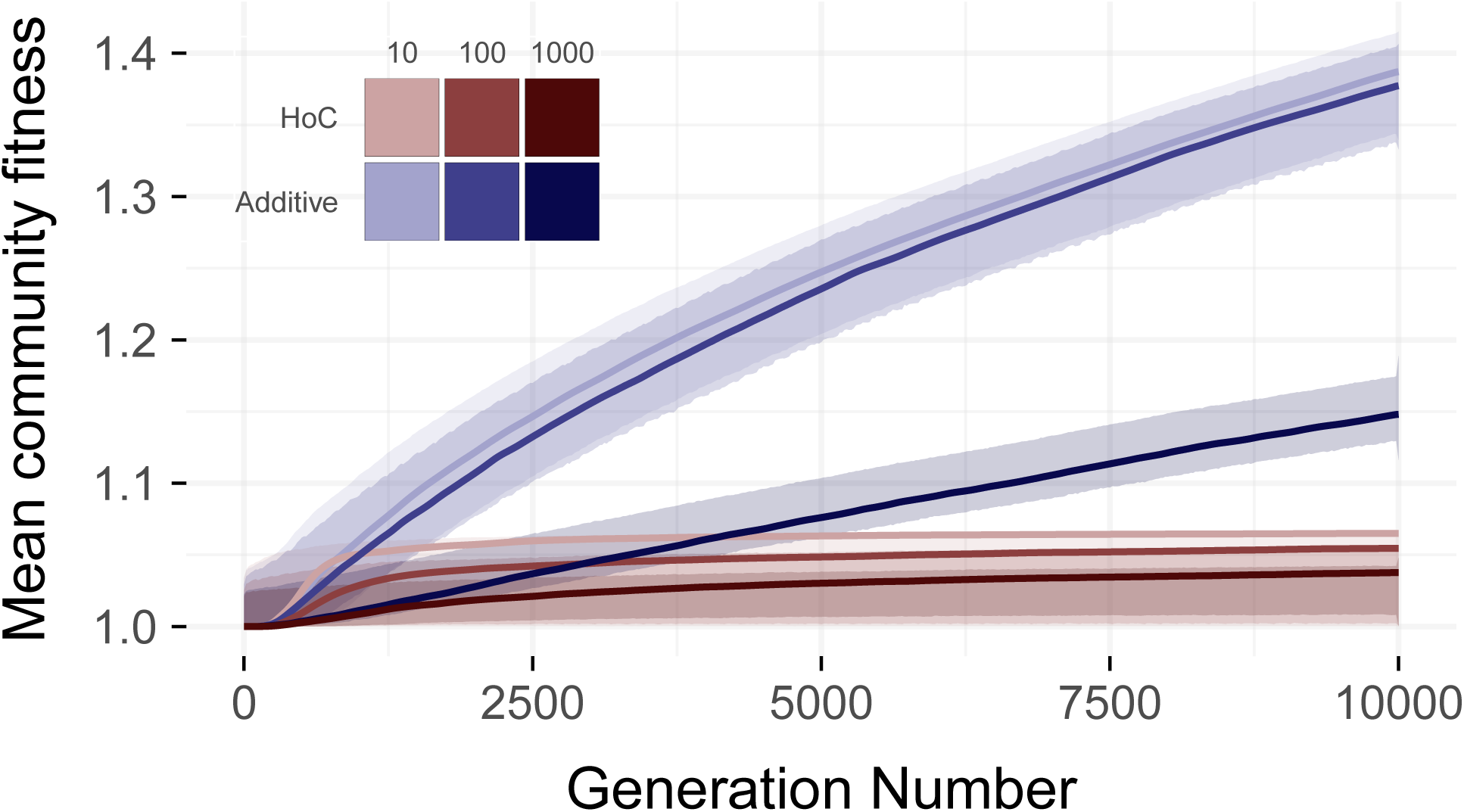
Community fitness through time simulated with rSHAPE. Colours represent results from different fitness landscape models (red - additive, blue - House of Cards, HoC) and shading reflects intensity of population bottlenecks. Solid lines represent the trend in mean community fitness across all replicates while the surrounding shaded polygon represent the absolute minimum and maximum values observed.

**Figure 3.**
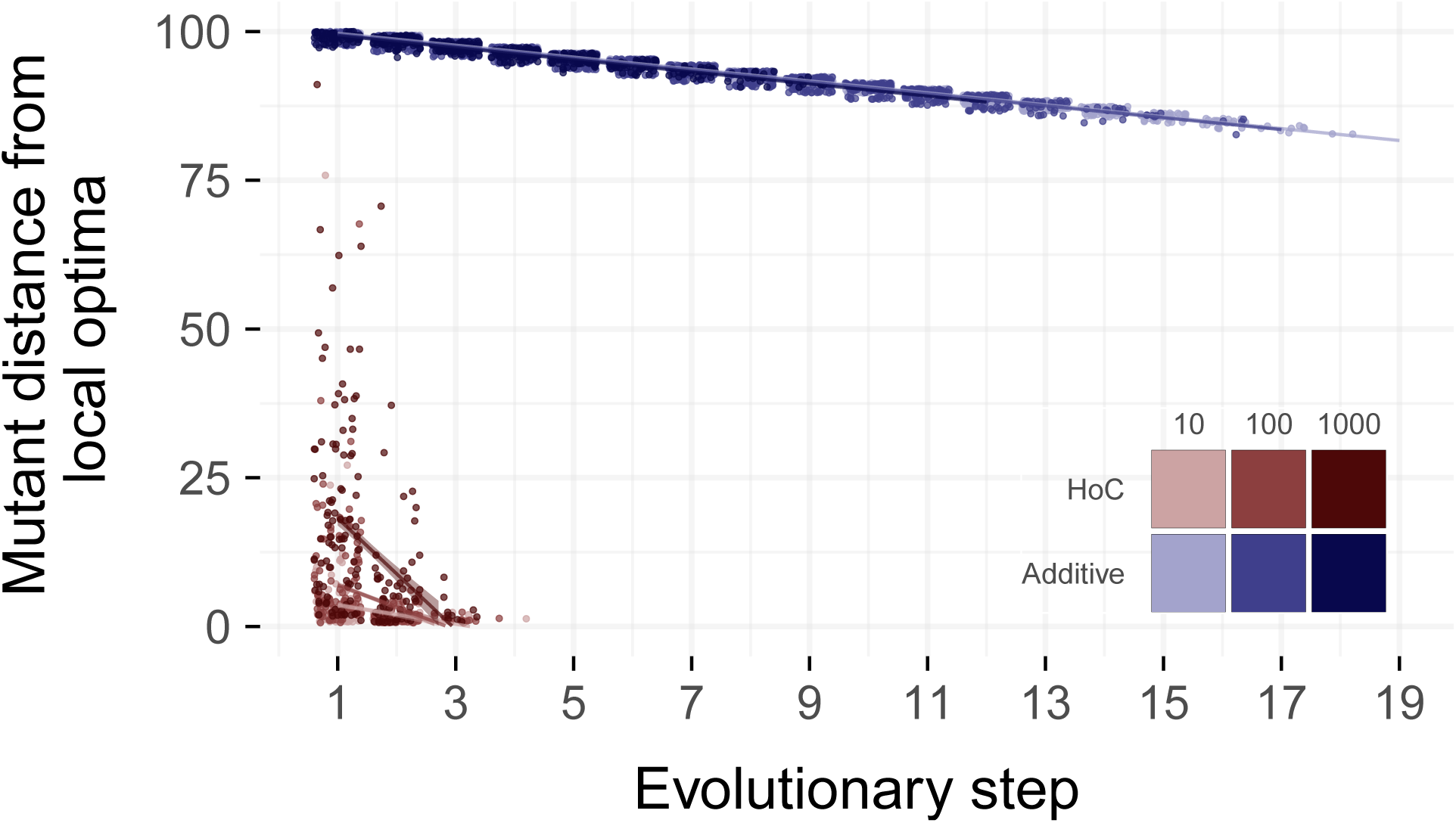
Distance of successful mutants from local optima as a function of ordered evolutionary step. Colours represent results from different fitness landscape models (red - additive, blue - House of Cards, HoC) and shading reflects intensity of regular population disturbance events (*i.e.* bottlenecks). Solid lines represent the average trend of mutational distance between “next evolutionary step” mutants and a local fitness optimum. Mutational distance represents the number of mutations separating two genotypes.

As my second assessment, I tested if rSHAPE could replicate seminal theoretical works related to the fixation probability of *de novo* mutants. The theoretical works compared herein all build upon the mathematical framework of Haldane (1927) who studied fixation probability in the simplest scenario of a constant size population. For all but the simplest scenario studied by Haldane (1927), I compared rSHAPE to the published approximate solutions (requiring simplifying assumptions) because the theoretical models of Ewens (1967); Otto and Whitlock (1997); Wahl and Gerrish (2001) would not otherwise have closed form solutions. All approximate solutions included at least the assumptions that selection coefficients (*s*) were “small” and that effective population sizes (*N*_*e*_) were “large” (no explicit values/formula were published for either). When comparing rSHAPE to these theoretical works, I simulated large populations with not less than one million individuals – a value that is within the range of population sizes considered by the works compared here. Since the exact fixation probability of Haldane’s model can be calculated, and because Haldane’s simplifying 2*s* approximation is part of all other theoretical works studied, I compared these two values to benchmark what may be considered a “small” selection coefficient. From this, I find that the approximate solutions may overestimate fixation probability by at least 5% once *s* ≥ 0.03 (Fig. 4B).

**Figure 4.**
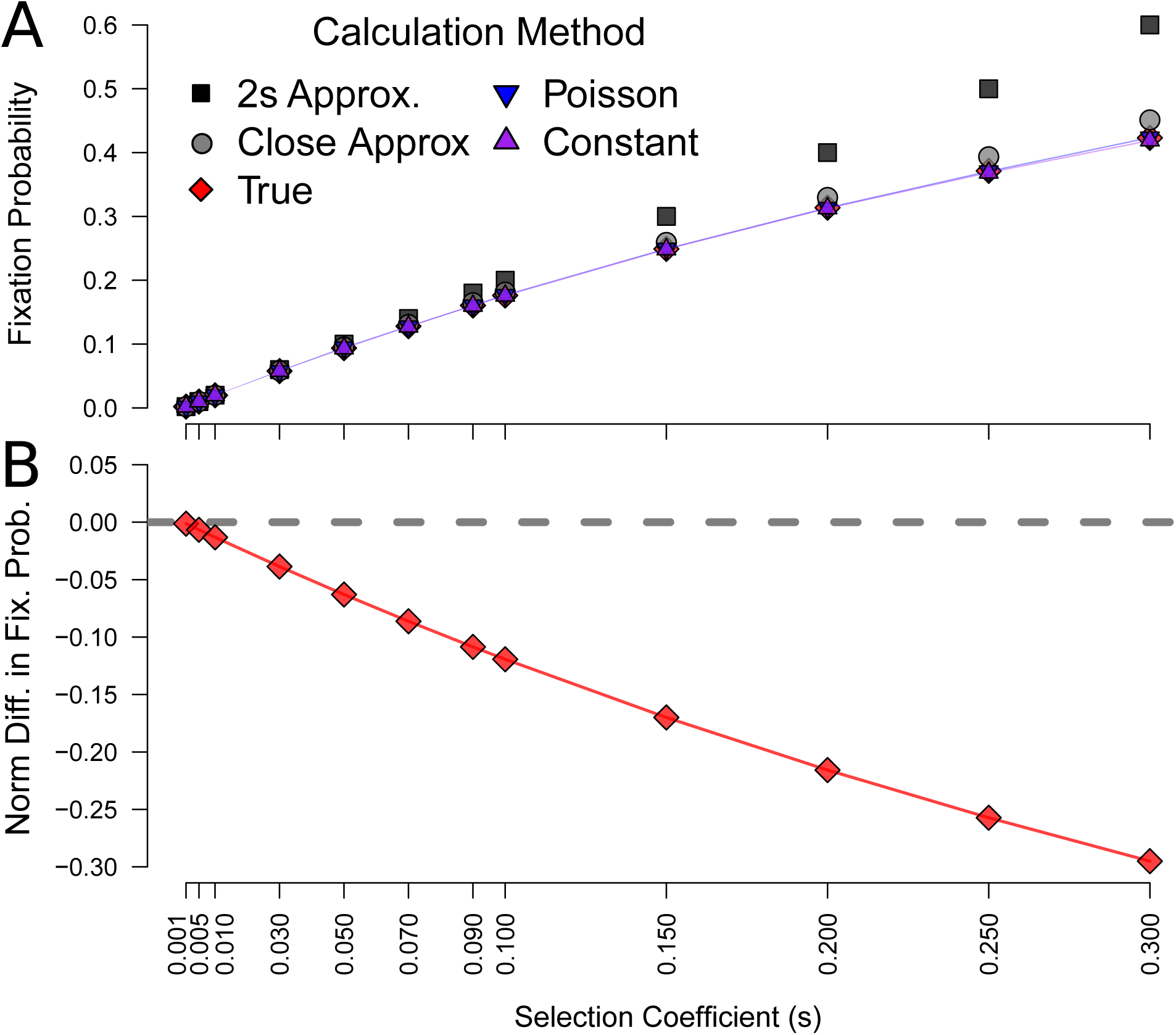
Theoretical and simulated probability of a single mutant fixing in populations of constant size. A shows the fixation probability, dependent on the selection coefficient *s*, where colouring of points distinguish between the theoretical calculations or simulated growth model. Theoretical calculations: black squares - *Haldane’s 2s* approximation, grey circles - the closer approximation 1 − *e*^−2*s*^; rSHAPE growth models: blue triangles - *Poisson*, purple triangles - *Constant* and; red diamonds represent the true un-approximated fixation probability. Note, the red diamonds are not always evident as they are overlain by the fixation probabilities estimated with rSHAPE. B, the normalised difference between the true fixation probability and the analytical approximation of 2*s*. Negative values reflect that the true fixation probability is lower than the analytical approximation of 2*s*.

As I compared rSHAPE to approximate solutions, I expected it to effectively replicate the-oretical fixation probabilities when model assumptions were not violated. From tests, I found that rSHAPE replicated the predicted fixation probability of *de novo* mutants arising in communities of constant size (Haldane, 1927) (Fig. 4, and Appendix **??**: “Haldane’s 2*s*”) as well as for those experiencing logistic growth (Ewens, 1967; Otto and Whitlock, 1997) (Fig. 5, and Appendix **??**: “Logistic Growth”). Under parameterisations which violated the underlying assumptions (*i.e.* high growth rate and/or large selection coefficient), rSHAPE underestimates fixation probability by an amount proportional to the difference observed when benchmarking “small” *s* (Figs. 4B and 5B). Thus, I have found that rSHAPE can replicate theoretically expected evolutionary dynamics under proper parameterisations. Further, from testing parameterisations that violated model assumptions, I found that rSHAPE predicts values that deviate from approximate solutions by an amount similar to the expected difference (from benchmarking “small” *s*) between the exact and approximate solutions of the underlying model. This suggests that rSHAPE provides a means to test evolutionary scenarios under conditions not supported by at least some analytical models.

**Figure 5.**
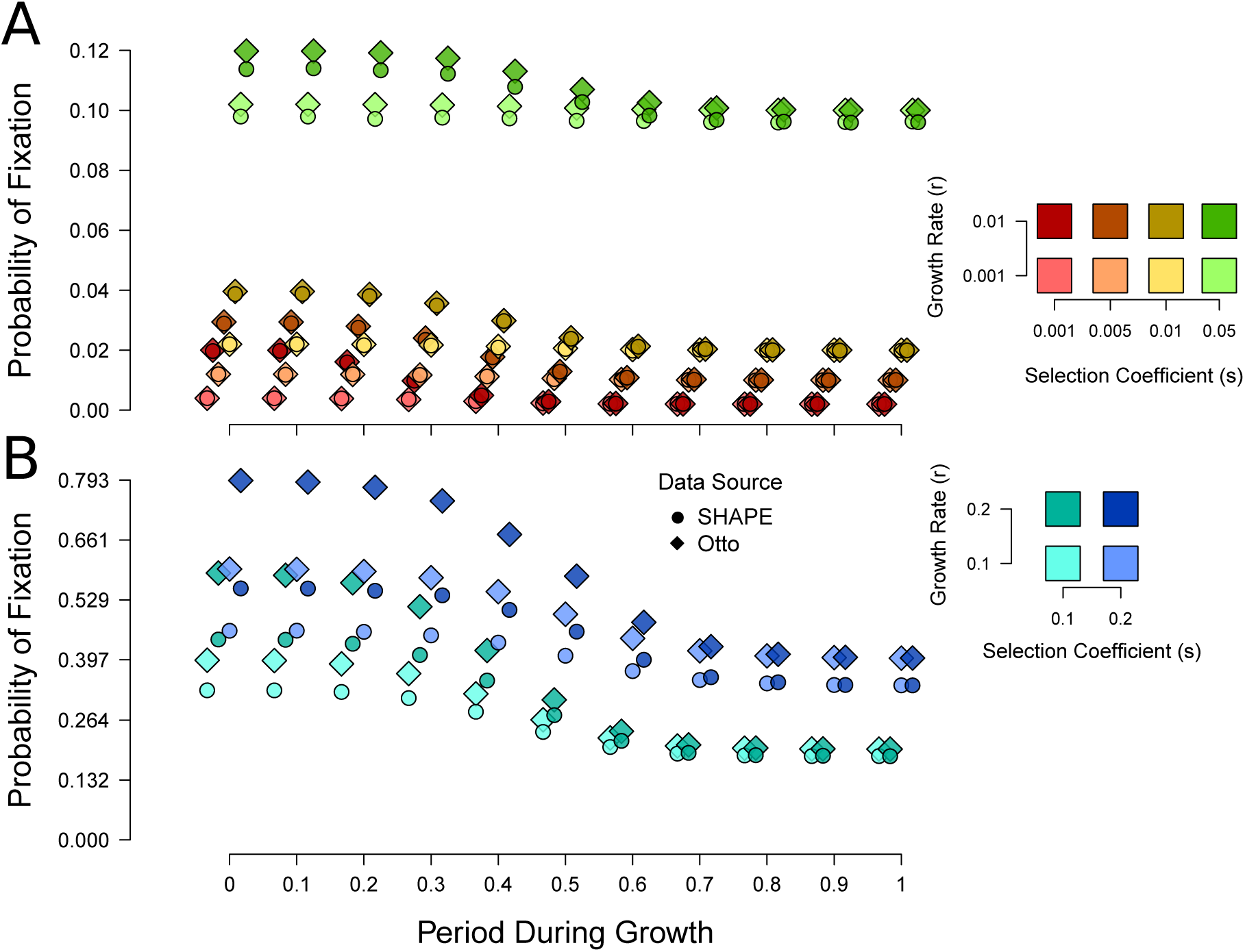
Theoretical and simulated probability of a single mutant fixing in populations growing to carrying capacity. Diamond shapes represent the analytical approximation of Otto and Whitlock (1997) while circles are for estimates using rSHAPE. The colour reflects the selection coefficient (*s*) with darker colours represent higher intrinsic growth rates (*r*). The period during the growth phase is scaled from the start of growth until the point where the number of individuals reaches carrying capacity. A presents the range of parameters originally studied in the theoretical work whereas B shows larger parameter values that implicitly violate assumptions of the analytical approximation.

All simulation scripts, and resulting summary files, can be found at github.com/Jdench/ rSHAPE_validation. The full results for this section are available upon request.

### rSHAPE application

Having validated the implementation of rSHAPE, I applied the simulation framework to assess the fixation probability of *de novo* mutants arising in a population that undergoes logistic growth and periodic population bottlenecks. It is common for microbial experimental evolution to include serial passaging, a practice that allows continued evolution by transferring a portion of stationary phase communities to fresh growth media. The process of serial passaging introduces periodic bottlenecks which delineate growth phases of the community and is expected to bias the phenotype of successful mutations (Wahl and Zhu, 2015). Earlier theoretical work suggests that the probability of a mutation both arising within the population and surviving repeated rounds of serial passaging is roughly uniform throughout growth phases that are effectively exponential (Wahl *et al.*, 2002). However, there are no estimates of how serial passaging affects fixation probability when growth is logistic in these communities.

To estimate this, I used rSHAPE to simulate the evolution of populations undergoing repeated rounds of serial passaging between growth phases that were modelled as either exponential or logistic growth. To compare the results of rSHAPE with previous theoretical work, I parameterised my simulations following the exponential and nutrient limited growth conditions of Wahl *et al.* (2002). The nutrient limited growth parameters were applied to my simulations of logistic growth. I found that rSHAPE does estimate a nearly uniform survival probability for *de novo* mutants arising in an exponential growth population (Fig. 6A). The parameters used in these simulations included a growth rate (*r* = 2) and mutation selection coefficient (*s* = 0.1) that would be deemed as “high” values outside parameter range of the simplifying assumptions in antecedent theoretical works (Haldane, 1927; Otto and Whitlock, 1997). Similar to my findings during validation, and because of the “high” selection coefficient and growth rate parameters, I had expected rSHAPE to estimate a lower fixation probability than the analytical approximation of Wahl *et al.* (2002), which it did (Fig. 6A). For conditions of logistic growth, I found that the joint probability of a mutant arising during the growth phase and subsequently surviving repeated rounds of serial passaging was not uniform (Fig. 6B). To better understand what evolutionary factors contributed to this result, I visualised and compared the individual probabilities that the mutant lineage arose during the growth phase, ultimately survived serial passaging, and also measured the expected mutant lineage size by the end of the growth phase in which it arose (Fig. 6C,D,E respectively). Note that the joint probabilities estimated for exponential and logistic growth differ by two orders of magnitude (Fig. 6A,B), this reflects the magnitude difference between the product of the initial populations size and mutation rate parameters used (*N*_0_ ∈ (10^5^, 10^7^) and *µ* ∈ (5 × 10^−5^, 4 × 10^−9^) for the exponential and logistic growth conditions respectively). This order of magnitude difference is reflected in the estimated probability of the mutant’s arrival throughout a growth phase (Fig. 6C). Under logistic growth, *de novo* mutants have a relatively higher probability of arising at all times up until the environmental carrying capacity is nearly reached (Fig. 6C). Yet, the probability of surviving repeated rounds of serial passaging is lower when growth is logistic (Fig. 6D), an observation likely underlain by the relatively smaller mutant lineage population size achieved in the first logistic growth phase (Fig. 6E).

**Figure 6.**
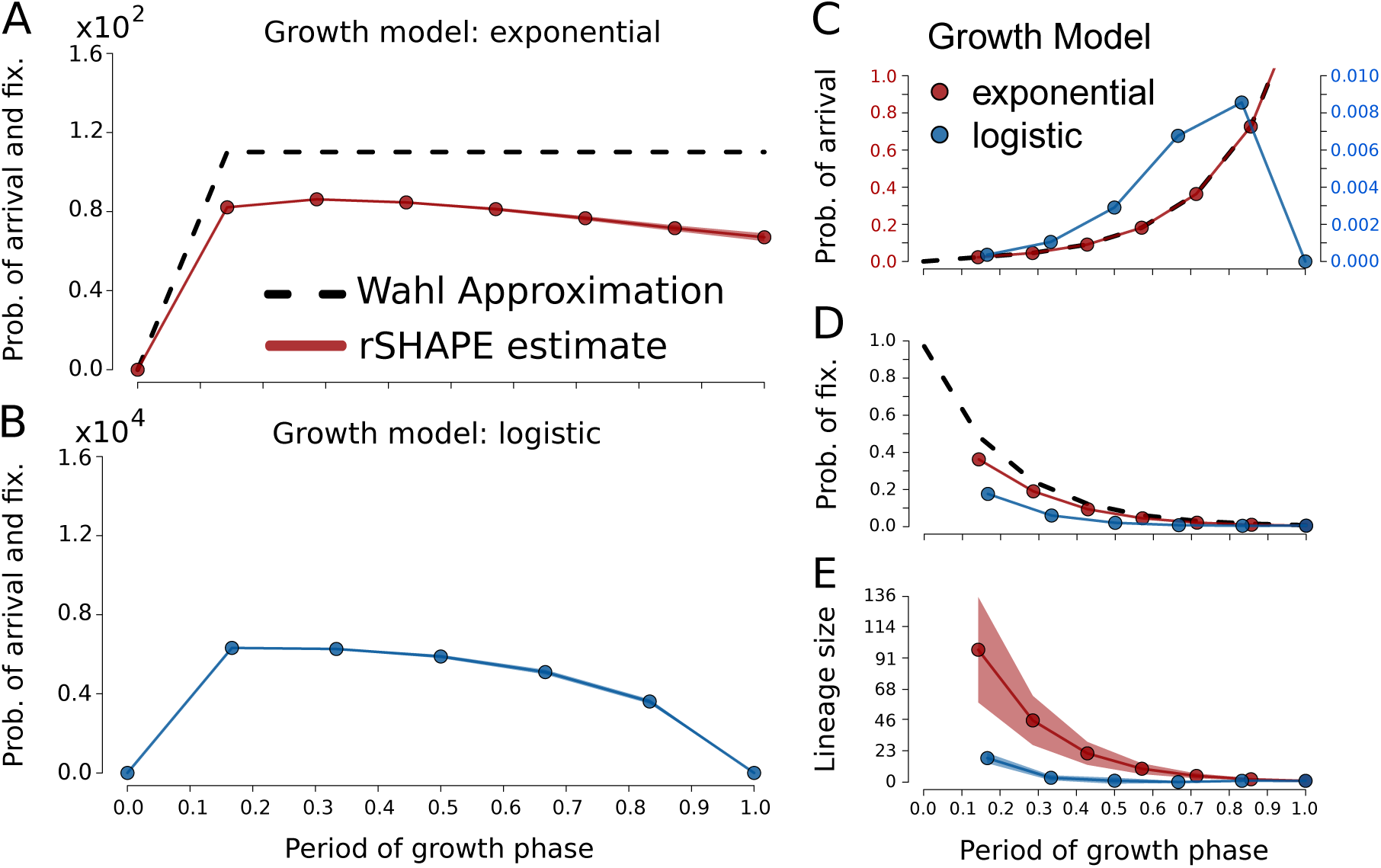
The joint probability of a single mutant arising during growth and then surviving many population bottlenecks, as a function of the time when a mutant arises during growth phases. Mutant selection coefficient *s* = 0.1, bottleneck strength was set to *D* = 100, and other parameters varied based on the model of growth (see main text). A compares the discrete analytical approximations of Wahl *et al.* (2002) to the estimates of rSHAPE for exponential growth. B shows the estimated probability of fixation when growth follows logistic form. C shows the independent probability of a mutant having arisen at a time during the growth phase whereas D visualises the probability to have ultimately survived repeated rounds of serial passaging. E shows estimates of the mutant lineage size by the end of the first growth phase in which it arose.

While a simple application, this example demonstrates how rSHAPE can be used to test hypotheses related to evolutionary scenarios where analytical modelling is not yet practicable. By applying rSHAPE to study the effects of serial passaging on *de novo* mutants arising during logistic growth, I provide evidence that the timing of mutant arrival during a growth phase does matter. If we assume that any, but not all, “next evolutionary step” mutants are equally likely to result from mutation events throughout the growth phase, then a biased fixation probability relative to time of arrival will increase stochasticity of survival for unestablished (*i.e.* newly arisen, small) mutant genotypes. Previous work has shown that microbial evolution experiments likely represent a case where an intermediate number of all possible mutants compete for fixation - a condition termed intermediate clonal interference (Bailey *et al.*, 2016). Intermediate clonal interference would suggest that only a subset of all mutants arise in any given generation and so serial passaging would introduce a “lottery” effect biasing fixation toward the subset of beneficial mutants arising first during an experiment. Since many microbial evolution experiments include serial passaging within their experimental design, it may be that serial passaging reduces the repeatability of genomic changes observed in those experiments. While this hypothesis remains to be tested, rSHAPE would be an ideal platform with which to perform such an analysis.

## Availability and future directions

rSHAPE is made available through CRAN or can be installed from GitHub github.com/JDench/ SHAPE_library. This software is made freely available under the GNU General Public License v3.0.

Future developmental priorities for rSHAPE are threefold. First, I would expand the applicability of rSHAPE by including methods to handle different habitat patches, partitioning of the genome, and temporal heterogeneity to selection. Second, implement additional fitness landscape models such Fisher’s Geometric Model(Fisher, 1958) and an eggbox model (Ferretti *et al.*, 2016). Lastly, I want to translate rSHAPE from the language of R to C in order to improve runtime through use of POSIX thread libraries and time for database queries.

## Acknowledgments

I would like to thank the Center for Advanced Computing (CAC) for access to their computational resources. I would also like to thank the Natural Sciences and Engineering Research Council (NSERC) of Canada for funding support accessed through the grants of my supervisors Drs. Stéphane Aris-Brosou and Rees Kassen.

## Notes

### Competing Interest Statement

The authors have declared no competing interest.

### Summary of Updates

Updated the text in preparation for submission to journal.

