## Supporting Information 1 for "The SHAPE of logistic growth shows that serial passaging biases fixation probability"

### Detailed overview of parameters and main functions

The framework of rSHAPE was written for, and implemented in, R R Core Team (2016). Nearly all functions imported by rSHAPE are part of the base R package. The exceptions being the SQL interface provided by *DBI* R Special Interest Group on Databases (R-SIG-DB) *et al.* (2016) and *RSQLite* Müller *et al.* (2017), the probability generating functions provided by *evd* Stephenson (2002), *VGAM* Yee and Wild (1996) and *sn* Azzalini (2016), the parallelisation functions of *foreach* Microsoft and Weston (2019), and the data concatenation function of *abind* Plate and Heiberger (2016).

#### General Parameters

The parameters described here are not an exhaustive list but instead those most likely to be of general interest for an rSHAPE user. It is further worth noting that some parameter combinations are intentionally redundant. This overparameterisation was intentional to facilitate the implementation of different models in rSHAPE.

A key parameter for any evolution experiment is its duration which rSHAPE controls through the number of generations $T$ to be simulated. In each generation, rSHAPE sequentially calls the stochastic events function, and the functions for deaths, births, and mutation events. Each of the deaths, births, and mutation functions has an associated probability parameter ($P_{d,b,m}$ respectively) which controls the per generation probability of any individual having an associated event. If $P_b$ is anything but 1, the term "generation" loses the traditional biological meaning of an average period of time between the birth and reproduction of an individual with average fitness. For convenience I herein use the term generation and time step interchangeably to mean a discrete unit of time simulated by rSHAPE.

The deaths and births of a population sum to represent their growth and the current growth models implemented in rSHAPE can simulate constant, exponential or logistic growth models. In rSHAPE, growth of evolving populations is calculated with the growth function that is a wrapper for the death and birth functions. Prior to running rSHAPE, users will have defined a focal number of individuals $N_f$ which is interpreted differently by the context of the growth model. In rSHAPE there are two constant number of individual growth models, named *Poisson* and *Constant*, wherein the focal number of individuals defines the starting number of individuals. The *Poisson* model comes from the Galton-Watson branching process Kimmel and Axelrod (2002) and assumes $P_d = P_b$ and that the intrinsic growth rate $r$ is two. This model will result in the population having a roughly constant size over many replicates, but in any one replicate this number is likely to deviate throughout time ($T$). I developed the *Constant* model to strictly enforce a constant number of individuals. The *Constant* model calculates the per genotype proportional number of births using the *Poisson* model and then scales these values to sum to the number of deaths. Note that for constant growth models the stochastic events function is ignored. For exponential growth, $N_f$ is the starting number of individuals and is used as the target number of individuals resulting from stochastic loss events. Under logistic growth, $N_f$ represents the environmental carrying capacity (*i.e.* $K$).

Throughout a run of rSHAPE, mutants arise with probability $P_m$ which is multiplied by either the number of births in a time step or, as one theoretical paper suggests, the total number of individuals Desai and Fisher (2007). An individual's genome has a constant length $L$ of binary state sites where 0 is the wild-type (WT) state and 1 the mutant. Sites in the genome have no explicit meaning other than that each are similarly mutable genomic regions which may affect genotype fitness. Users may define if revertant mutations are permitted (*i.e.* if 1 can mutate to 0). The fitness $w_i$ of a genotype $i$ (where $i \in \{1, 2, 3, ..., x\}$ and $x$ is the number of unique living genotypes) is calculated given the fitness landscape model parameter that often depends upon a parameterised random effect distribution. The fitness landscape models currently implemented are: Additive, Fixed, House of Cards (HoC) Kingman (1978), Kauffman's NK (NK) Kauffman and Weinberger (1989), or Rough Mount Fuji (RMF) Aita *et al.* (2000); Neidhart *et al.* (2014). The additive model draws the effect (*i.e.* selective coefficient) of mutations from the random effect distribution and calculates $w_i$ as the sum of the mutation effects (*i.e.* no epistasis). The fixed model is only practical for small genotypes as it requires that the user supply a matrix defining the fitness value for each genotype, but it does allows users experiment with defined (*i.e.* fixed) fitness landscapes. A full description of the HoC, NK and RMF models are beyond the scope of this work but in brief each calculates $w_i$ using values drawn from the random effect distribution, and some require additional constants that can be defined in rSHAPE. At present, rSHAPE implements the following probability generating functions for use as the random effect distribution: Beta, $\chi^2$, Exponential, Frétchet, Gamma, General Extreme Value Distribution Stephenson (2002), Normal, Reverse Weibull, Skew Normal Azzalini (2016) and Uniform.

Once initial parameters are defined and a run starts, each generation will begin with a call to the population disturbance function that implements stochastic loss events.

#### Population Disturbance

This function is used to simulate disturbance events that stochastically reduce the total number of individuals. Disturbance events occur based on a schedule defined either by a fixed number of generations between events or where subsequent events are scheduled so that the expected number of births prior to the next event balances the sum of lost individuals. The expected number of individuals lost for each genotype $i$ is proportional to their frequency within the evolving population. Using the disturbance factor $D$, the new expected number of individuals of each genotype $i$ is calculated as $\frac{N_i}{D}$ . This process is stochastic because the actual number of remaining individuals is calculated as random draw from a Poisson distribution with location parameter $\frac{N_i}{D}$. For simulations using exponential growth, the value $D$ is controlled by $N_f$ such that:

$$D = \sum_{i=1}^{x} N_i / N_f \tag{1}$$

where $N_i$ is the number of individuals of genotype $i$. This prevents exponential population growth from growing indefinitely provided there are population disturbance events. As constant growth models ignore population disturbance events, the user can only meaningfully set $D$ when growth is logistic. When population growth is logistic, $D$ can either be a constant value or be randomly drawn from a normal distribution parameterised by the user. Any value $D < 1$, either as a constant or resulting from random draws, will automatically be set to one since this function should never increase population sizes.

Next rSHAPE will calculate growth of the evolving population.

#### Growth

In rSHAPE, deaths and births happen simultaneously meaning that in a single time step an individual may both die and produce offspring. However, deaths are calculated first in order to inform certain growth model calculations.

**Death Events**

The probability of death $P_d$ controls the proportion of individuals that die in a generation. The user can choose if this probability is a constant value or if it is applied in a density dependent manner. When $P_d$ is constant, the number of deaths for a genotype $i$ is given by:

$$deaths_i = N_i \, P_d \tag{2}$$

where $N_i$ is the number of individuals of genotype $i$. If deaths are density dependent the number is calculated by:

$$deaths_i = N_i \, P_d \, (\frac{\sum_{i=1}^{x} N_i}{K_d})^{c_d} \tag{3}$$

where $K_d$ is the population size at which $P_d$ is 100% its defined value. The exponent $c_d$ is used to scale the product of $P_D$ and the ratio of population size and $K_d$, where larger $c_d$ causes $P_d$ to have little impact until the population is close to $K_d$. Please note that $K_d$ is defined separately from the logistic growth carrying capacity $K$ (*i.e.* $N_f$).

This function is called prior to births but does not directly affect the population sizes ($N_i$) used in birth calculations. The number of deaths is first calculated so that births can be scaled to deaths such as under conditions of the *Constant* growth model.

**Birth Events**

The per generation probability of any individual giving birth is controlled by $P_b$. The number of offspring generated by an individual is calculated given the growth model parameterised by the intrinsic growth rate *r*, and the genotype's fitness $w_i$. For the *Constant* growth model, the number of births for each genotype is proportional to their size $N_i$ and is calculated in two steps. The first is to calculate the absolute birth potential of each population by

$$births_{potential_i} = N_i \, (1 + w_i - \bar{w}) \, P_b \tag{4}$$

where the term $(1 + w_i - \bar{w})$ calculates relative fitness centred around 1 using the mean fitness $\bar{w}$. This method of calculating relative fitness handles instances when $\bar{w} = 0$ but is sensitive to the magnitude of $\bar{w}$. The user can choose relative fitness to be calculated using the more traditional $\frac{w_i}{\bar{w}}$, but this will be overridden if $\bar{w} = 0$. The second step in calculating the *Constant* growth uses the potential births ($births_{potential_i}$) calculated in eq. 4 as weightings in the final calculation:

$$births_i = \frac{births_{potential_i}}{\sum_{i=1}^{x} births_{potential_i}} \sum_{i=1}^{x} deaths_i \tag{5}$$

where the actual number of births is $births_{potential_i}$ scaled to the sum of deaths. If no potential births occurred, or there were no deaths, then the second step is skipped and $births_i$ is returned as a vector of zeroes. If the *Poisson* growth model was chosen, then $births_i$ is obtained through draws from a Poisson distribution where the location parameter is given by the product of $N_i$, $w_i$, and $P_b$. This approach was derived from the theoretical work of *Haldane* (1927) and assumes a large population and that $P_b = P_d$.

To calculate births when growth is either exponential or logistic growth, then the intrinsic growth rate $r$ is an additional parameter controlling the number of births for each population. For exponential growth the expected number of births for each genotype are calculated as

$$births_i = N_i\,(e^{ln(r)\, w_i\, P_b} - 1) \tag{6}$$

where $ln(r)$ is the natural logarithm of the intrinsic growth rate and $w_i$ is the fitness of a genotype. This equation is derived from the exponential growth model but I subtract 1 from the growth term to result in a calculation for births. Note that if $e^{ln(r)\, w_i\, P_b}$ [illegible] 1 then the result it is set to zero to prevent population decrease being calculated by the birth function. When growth is logistic, the expected number of births for each genotype is calculated in two steps. First, a density dependent growth term $dg_i$ for each genotype $i$ is calculated using the logistic equation

$$dg_i = N_i + (w_i\, r\, P_b)\frac{K - \sum_i^x N_i}{K} \tag{7}$$

where $K$ represents the carrying capacity. This density dependent term $dg_i$ represents the amount of growth expected for genotype $i$ given the current total number of individuals. Using $dg_i$, the number of births for each genotype is calculated as

$$births_i = N_i\,(\frac{dg_i}{\sum_i^x N_i} - 1) \tag{8}$$

where similar as to with exponential growth, I subtract 1 to calculate births rather than growth. Recall that while deaths are calculated prior to births they do not affect $N_i$ used in these calculations. The exponential and logistic growth model birth calculations are deterministic and so to make growth calculated by rSHAPE a stochastic process, users can toggle that "drift" be considered. "Drift" causes the number of births to become random draws from a Poisson distribution using location parameters set by the deterministic birth values. Also, since births and deaths are calculated separately, but because users may want growth to perfectly simulate deterministic growth models, users may set the number of births calculated to be scaled by deaths. In this case, any deviations will be adjusted by generating additional births, or deaths, as required for each population using a nested call to the growth function with the *Constant* method.

Once growth has been calculated, rSHAPE will determine if mutants are generated.

#### Mutation Events

The last step of each generation is to calculate if there have been mutation events. The number of mutations is controlled principally by the per genome, per generation, mutation rate $\mu$. Classically the number of mutants is a product of $\mu$ and the number of replication (*i.e.* birth) events, but more recent theoretical work has suggested using all living individuals Desai and Fisher (2007). Either choice may be simulated with rSHAPE by selection of a logical toggle parameter. The number of mutants generated from replication events is calculated as:

$$mutants_i = \mu\, births_i\, \frac{r}{r-1} \tag{9}$$

where recall $r$ is the number of offspring expected from a single birth event and the term $\frac{r}{r-1}$

ensures that the number of mutants considers not just offspring but also the parental individuals since either may mutate. If mutants can arise from any individual in the population, rSHAPE adds the following vector:

$$\mu(N_i - deaths_i - \frac{births_i}{r-1}) \tag{10}$$

to the values calculated in eq. 9. The first time any genotype generates mutants, rSHAPE will calculate, and permanently record, the fitness value of all genotypes in the unexplored neighbouring mutational space. The fitness for a genotype is calculated based on the chosen fitness landscape model (see *General Parameters* above). For each genotype $i$ with at least one mutant, rSHAPE draws $mutant_i$ times (with replacement) from the list of genotypes in the neighbouring mutational space.

### Detailed comparisson to theoretical work

#### Haldane's $2s$

The seminal theoretical work of Haldane Haldane (1927) approximates the probability of fixation for a single mutant as being $\approx 2s$ (where $s$ is the selection coefficient). Haldane's work makes the assumption that the WT population is very large, that the number of individuals is constant, that $s$ is small ($s \ll 1$), that generations do not overlap, and that a successful mutant lineage must grow from a single progenitor mutant. I replicated these conditions in rSHAPE with each of the two appropriate growth models: *Poisson* and *Constant*. The first model is identical to the method used by Haldane Haldane (1927) and is called the *Poisson* form because it is based on a Galton-Watson branching process Kimmel and Axelrod (2002) and estimates birth events as draws from a Poisson distribution. This method of simulating births causes a populations size to be approximately constant over many replicates but allows population size to vary within any one replicate. In microbial experimental evolution, growth conditions can lead to population sizes being roughly constant by carefully balancing nutrient flux ($e.g.$ chemostat grow chamber). The *Constant* growth form better simulates these conditions by scaling the births of the *Poisson* form to be exactly the number of deaths in a generation.

Using a range of selective coefficients, $s \in \{0.001, 0.005, 0.01, 0.03, ..., 0.09, 0.1, 0.15, ..., 0.3\}$, I calculated the fixation probability from 1,000,000 replicate runs of rSHAPE. Haldane's $2s$ approximation comes from eq. 11 (eq. 1 of Haldane's theoretical work Haldane (1927))

$$s = \sum_{i=2}^{\infty} \frac{prob_{fix}^{i-1}}{i} \tag{11}$$

where $prob_{fix}$ is the probability of fixation. The classic approximation $prob_{fix} \approx 2s$ considers only the first term of eq. 11 however, the analytical expression of $1 - e^{-2s}$ provides a closer approximation (personal communications with Dr. Lindi Wahl). We can calculate the exact value for $prob_{fix}$ related to $s$ (up to finite level of precision) by exploring the space of $prob_{fix}$ values and plugging them into the rearranged, and un-simplified form, of eq. 1 from Haldane's work:

$$s = \frac{-ln(1 - prob_{fix})}{prob_{fix}} - 1 \tag{12}$$

the search space should be centered around $2s$ and can continue to adjust by $prob_{fix} \pm \delta x$ until $s$ is found. I found that both the *Poisson* and *Constant* growth forms implemented in rSHAPE accurately predict exact fixation probabilities calculated in this way (Fig. **??**A, main text). As $s$ increases, both the close approximation and $2s$ would suggest higher fixation probabilities. One of the assumptions of Haldane's work is that $s$ is small. As "small" is a subjective term, I compared $2s$ against the true fixation probability finding that $2s$ overestimated fixation probability by $\sim 5\%$ when $s = 0.05$ and that this difference increased with $s$ (Fig. **??**B, main text). In fact only when $s < 0.01$ does the $prob_{fix}$ approximation of $2s$ differ by less than 1.3% from the exact value. This result suggests that $s$ be considered small, for theory building upon Haldane's work, only when $s \leq 0.01$.

The work described here was for a large population ($N_f = 10^8$), but I did also run simulations with a smaller population ($N_f = 10^4$) and found that the absolute probability of fixation values were still quite in line with the exact values (Fig. 1A). However, the normalised difference between the exact and estimated probability of fixation values differed significantly, for the small population only, when $s \leq 0.03$ (Fig. 1B).

#### Logistic growth

Microbes grown experimentally tend to follow a logistic growth pattern with an upper bound (*i.e.* carrying capacity $K$) caused either by density dependent death or reduction in birth rate such as when nutrients are depleted. Theory suggests that the fixation probability of a single *de novo* mutant is greater than $2s$ relative to the growth rate of the population and becomes $\sim 2s$ once $K$ is reached Ewens (1967); Otto and Whitlock (1997). Implicitly, this assumes that while the community is at $K$ births continue but are balanced by deaths. In my simulations replicating the work of Otto and Whitlock (1997), the assumptions are all similar to those of Haldane's $2s$ but, the growth rate $r$ is assumed to be small. I performed simulations with the same range of $s$ as previously described and that range was also used for $r$. Populations were initiated at $\frac{K}{100}$ and I simulated growth for two carrying capacities $K \in 10^{(4,8)}$ but only discuss the results for the larger as both are similar (Fig. 2). To calculate the theoretical fixation probability, I used the analytical approximation shown in eq. 13 (eq. 11 from Otto and Whitlock (1997))

$$prob_{fix}(t) \approx \frac{2sK(s+r)}{sK + rN_{WT}(t)} \tag{13}$$

where $prob_{fix}(t)$ is the fixation probability given the mutant arose at time $t$ when there were $N$ wild-type individuals in an environment with carrying capacity $K$. Estimates with rSHAPE were calculated from 1,000,000 replicates.

When both $s$ and $r$ are within the range of values presented in the work of Otto and Whitlock Otto and Whitlock (1997), rSHAPE accurately reproduces the analytical approximation except once either is equal to or greater than 0.05 where rSHAPE estimates visibly lower fixation probabilities (Fig. S2). Similar to my findings when comparing $2s$, I find that rSHAPE underestimates the probability of fixation by an amount proportional to the value of either $r$ or $s$ when they are not "small" (*i.e.* the assumptions are violated).

#### Serial Passaging

The laboratory practice of serial passaging, common to microbial experimental evolution, induces regular population bottlenecks as a small proportion of a community grown in liquid media is transferred to new media for continued growth. The ratio of new to old volume is commonly expressed as the dilution factor $D$ and the period between transfers is known as the growth phase. Prior to transfer, the old media should be well mixed so that the existing genotypes are proportionally transferred. However, these repeated bottlenecks result in rare genotypes being lost Wahl *et al.* (2002) because in practice sampling is still a random process and even when samples are perfectly mixed we are not likely to transfer any genotype with less than $D$ individuals. Through the stochastic events function, rSHAPE can simulate serial passaging which permitted this last comparison. Current theory has suggested that the probability of a mutant arising and eventually fixing is roughly uniform throughout the growth phase when a population grows exponentially Wahl *et al.* (2002). After validating that rSHAPE accurately simulates serial passaging (Fig. S3), I used it to estimate the combined probability that a mutant will arise at some point throughout a growth phase and then eventually fix.

To make the maths tractable, the work of Wahl *et al.* (2002) assumed that growth was effectively exponential, that there was no competition and that deaths could be ignored. I ran simulations using these conditions and the same parameters from the exponential and nutrient limited growth models of Wahl *et al.* (2002). The later conditions were applied to logistic growth in rSHAPE.

For the exponential (logistic growth) models, I began independent simulations by seeding a single mutant ($s = 0.1$) into a population of $10^5(10^7)$ individuals at each of the $\tau = 7$ (6) generations in the growth phase. The basal growth rate was $r = ln(2)$ ($r = 2$) and each run ended when the mutant was lost, had fixed, or one million serial passaging events of $D = 100$ had occurred at which point fixation was assumed imminent. To estimate the fixation probability, each combination of parameters was replicated 1,000,000 times. I compared estimates against the analytical approximation for survival (eq. 15) which depends upon the extinction probability ($P_{extinction}$) given by eq. 14 Wahl and Gerrish (2001); Wahl *et al.* (2002)

$$P_{extinction} \approx 1 - 2se^{-rt}r\tau \tag{14}$$

where $t$ is the generation of the growth phase during which the mutant arises. The probability of a mutant with selective coefficient $s = 0.1$ having been born at time $t$ was calculated as the product between the mutation rate $\mu = 5 \times 10^{-5}(4 \times 10^{-9})$, the number of births in a generation, and the probability density of an exponential distribution with rate $\alpha = 100$ (value was not published in Wahl *et al.* (2002) and is as per personal communications with Dr. Wahl) and subsequently survives. So, the analytical approximation of a mutant arising during the growth phase and ultimately surviving multiple serial passaging events was calculated with eq. 15 Wahl *et al.* (2002)

$$P_{survival} \approx births(t)\, \mu\alpha e^{-\alpha s}(1 - P_{extinction}) \tag{15}$$

where the number of births is the difference in population size between generations $t$ and $t-1$ (because deaths are ignored). When growth is exponential, rSHAPE estimates a roughly uniform joint probability throughout the growth phase though the exact value is lower than the analytical approximation. This lower estimate is not surprising as the theoretical work being compared is built upon Haldane's $2s$ approximation.

When growth was logistic, rSHAPE estimated a joint probability that is not uniform. This result suggests, similar to other theoretical work Wahl and Zhu (2015), that the protocol of serial passaging can affect which mutations are fixed during an experiment. Because microbial populations used in experimental evolution tend to grow logistically, the practice of serial passaging will bias the fixation of mutations arising early during growth. Early in the growth phase there are fewer mutants being generated which means the mutational space will be less well explored and so of those mutants which arise and fix may be lottery winners instead of mutationally optimal alternatives. It could be argued that if the time to fixation were long then we might expect the dynamics to replicate maximal clonal interference as all mutants eventually arise multiple times and compete for fixation. Under maximal clonal interference we expect mutationally optimal outcomes as selection coefficients drive evolution. However, previous work suggests that the dynamics of microbial experimental evolution reflect intermediate clonal interference Bailey *et al.* (2016) whereby some, but not all, possible mutants compete for fixation and so the order in which they appear matters (*e.g.* timing during growth phase). Other studies have shown how the order in which mutations appear can affect evolutionary outcome Weinreich *et al.* (2005); Kvitek and Sherlock (2011); Sackman and Rokyta (2017). With rSHAPE I have shown that the practice of serial passaging introduces a jackpot scenario where those few mutants arising early during growth phases are most likely to survive. This finding suggests that the stochasticity in outcome between replicate microbial experimental evolution populations is at least somewhat due to the practice of serial passaging. Future studies should use rSHAPE to quantify what proportion of stochasticity is attributable to serial passaging.

R Core Team 2016. *R: A Language and Environment for Statistical Computing*. R Foundation for Statistical Computing, Vienna, Austria.

R Special Interest Group on Databases (R-SIG-DB), Wickham, H., and Müller, K. 2016. *DBI: R Database Interface*. R package version 0.5-1.

Sackman, A. M. and Rokyta, D. R. 2017. Additive phenotypes underlie epistasis of fitness effects. *Genetics*.

Stephenson, A. G. 2002. evd: Extreme value distributions. *R News*, 2(2): 0.

Wahl, L. M. and Gerrish, P. J. 2001. The probability that beneficial mutations are lost in populations with periodic bottlenecks. *Evolution*, 55(12): 2606–2610.

Wahl, L. M. and Zhu, A. D. 2015. Survival probability of beneficial mutations in bacterial batch culture. *Genetics*, 200(1): 309–320.

Wahl, L. M., Gerrish, P. J., and Saika-Voivod, I. 2002. Evaluating the impact of population bottlenecks in experimental evolution. *Genetics*, 162(2): 961–971.

Weinreich, D. M., Watson, R. A., and Chao, L. 2005. Perspective: Sign epistasis and genetic constraint on evolutionary trajectories.

Yee, T. W. and Wild, C. J. 1996. Vector generalized additive models. *Journal of Royal Statistical Society, Series B*, 58(3): 481–493.

### Figures

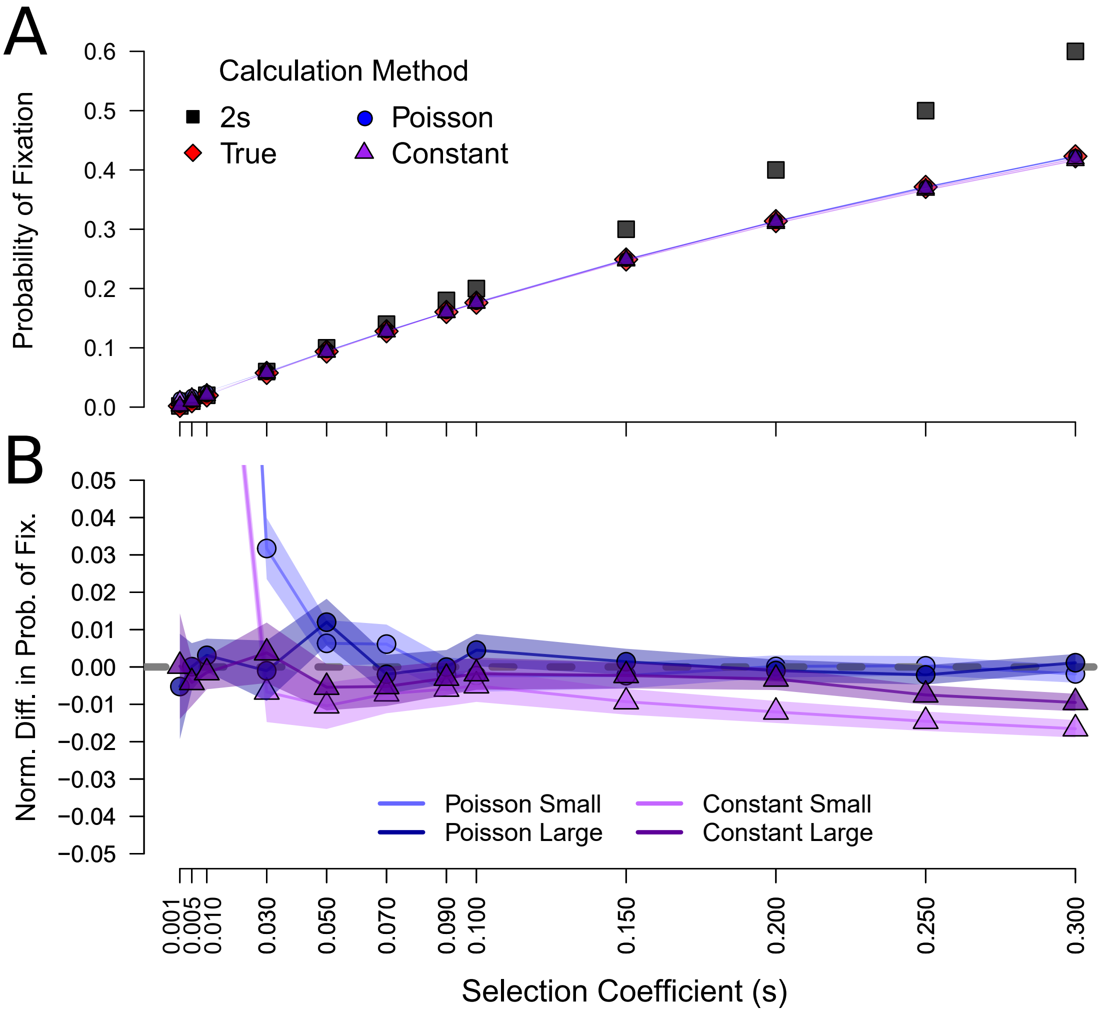


**Figure 1.** **Comparison between the theoretical and estimated probability of a single mutant fixing, in an environment with a constant number of individuals.** The colour and shape of points identify the means for calculating/estimating fixation probability whereas the shading identifies if simulated estimates were performed with smaller ($10^4$) or larger ($10^8$) population sizes. Panel A shows the fixation probability, dependent on the selection coefficient $s$, and the red diamonds show the un-approximated theoretical expectation (*i.e.* true fixation probability). Panel B presents the normalised difference between the true fixation probability and the values calculated with rSHAPE. Negative values reflect when the values calculated with rSHAPE are lower than the true fixation probability.

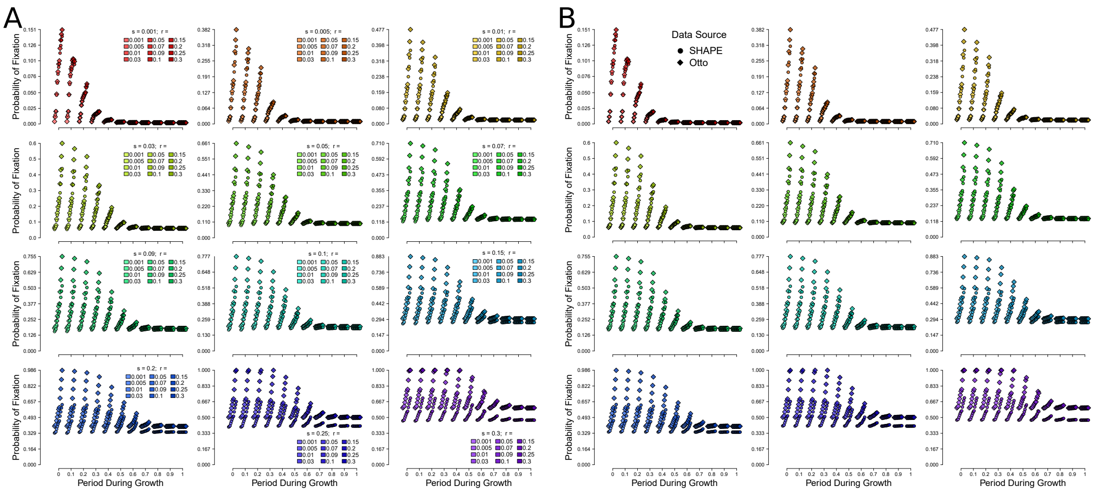


**Figure 2. Comparison of the theoretical and estimated fixation probability for a mutant growing logistically.** Diamond shapes represent the analytical approximation of Otto and Whitlock (1997) while circles are for estimates using rSHAPE. The colour used to fill points reflects the selection coefficient $s$ and darker colours represent higher intrinsic growth rates $r$. The period during the growth phase is scaled from the start of growth until the point where the number of individuals reaches carrying capacity. The range of parameters is similar between panels but A is for when $K = 10^4$ whereas B is for $K = 10^8$.

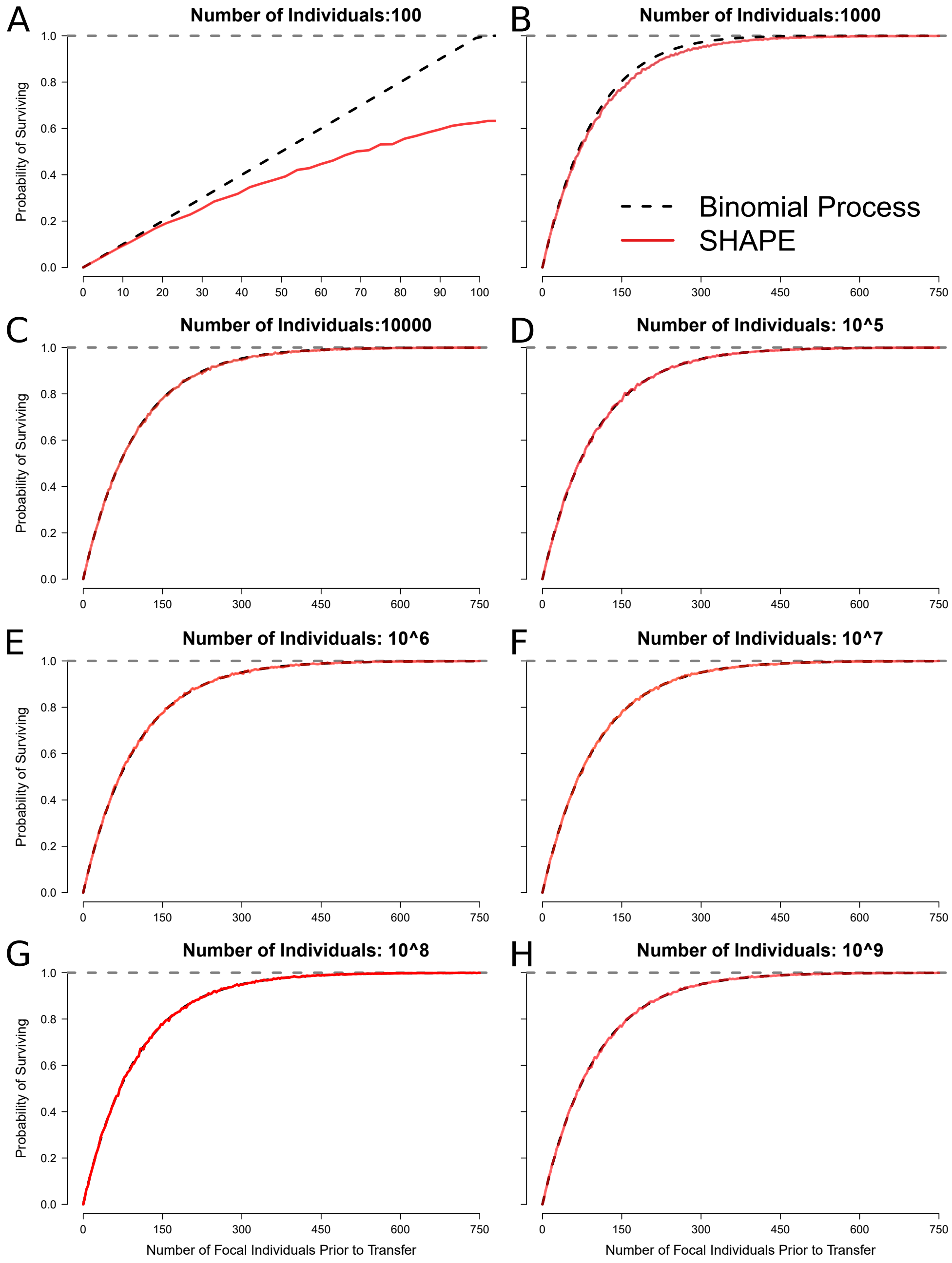


**Figure 3.** **The probability that at least one individual of a focal genotype survives a disturbance event (serial passaging) reducing the total number of individuals 100 fold ($D = 100$).** The red polygon (appears as a line) shows the 95% CI of survival probability calculated with rSHAPE, while the black dashed line shows the expectation calculated from a binomial process. The dashed grey line highlights when probability reaches unity. Panels differ in the number of focal individuals, and total population size, prior to transfer.

### Tables

**Table 1**

**List of definable evolutionary parameters in rSHAPE.** While rSHAPE offers a variety of modifiable evolutionary parameters, all have default values and many interact with others as part of calculations (*e.g.* The probability of birth ($P_b$) and intrinsic growth rate ($r$) will both affect the number of offspring calculated). Where possible, default values were chosen to best reflect standard parameterisation of evolutionary models involving haploid asexuals. For more details, please refer to the reference material included with rSHAPE.

| Context | Symbol | Parameter & Meaning |
|---|---|---|
| Individual | $L$ | The number of genome positions. |
| Disturbance | | Type of disturbance; either fixed or random bottlenecks. |
| Events | $D$ | Dilution factor of the first disturbance event. |
| | | Value(s) used to calculate size of disturbance events. |
| | | Number of time steps between disturbance events. |
| Birth | $P_b$ | Probability that an individual produces offspring in a time step. |
| | | Growth model to be used. |
| | $r$ | Number of individuals after a birth event (*i.e.* : parent + offspring) |
| | | Logical toggle to add randomness to birth calculations. |
| | $N_f$ , $K$ | Focal population size used conditionally on growth model. |
| Death | $P_d$ | Probability that an individual dies in a time step. |
| | | Logical toggle controlling if deaths density dependent. |
| | $c_d$ | Parameter controlling shape of density dependence. |
| | $K_d$ | Population size at which 100% of density dependent deaths occur. |
| | | Are births scaled to replace all deaths? Enforces adherence to growth model calculations. |
| Mutation | $P_m$ | Probability of a mutant arising during a time step. |
| | | Logical toggle: Is mutation probability only applied to individuals undergoing birth? |
| | | Can mutations revert to wild-type state? |
| Fitness | | Fitness landscape model used to calculate genotype fitness. |
| Landscape | | Distribution used to draw random component of fitness calculations. |
| | | Parameterisation of distribution used to draw random fitness components. |
| | | Fitness value of wild-type genotype. |
| | | Logical toggle: are genotype fitness calculations to be used as selection coefficients or relative fitness? |
| | | For RMF fitness landscape model: number of mutations separating the wild-type and optimal genotypes. |
| | | For RMF fitness landscape model: weighting of the independent fitness component. |
| | | For NK fitness landscape model: Number of sites interacting. |
| Experiment | $T$ | Number of discrete time steps to simulate. |
| | $n$ | Number of replicates for a parameter combination. |
| | | Proportion of a general time step calculated in each realised time step. |
| | | Logical toggle: are individuals tracked as integer values or allow decimal tracking. |
